# Nonuniform structural properties of wings confer sensing advantages

**DOI:** 10.1101/2022.10.04.510910

**Authors:** Alison I Weber, Mahnoush Babaei, Amanuel Mamo, Bingni W Brunton, Thomas L Daniel, Sarah Bergbreiter

## Abstract

Sensory feedback is essential to both animals and robotic systems for achieving coordinated, precise movements. Mechanosensory feedback, which provides information about body deformation and position in space, depends not only on the properties of sensors but also on the structure in which they are embedded. In insects, wing structure plays a particularly important role in flapping flight: in addition to generating aerodynamic forces, wings provide mechanosensory feedback necessary for guiding flight while undergoing dramatic deformations over the course of each wingbeat. However, the role that wing structure plays in determining mechanosensory information is relatively unexplored. Insect wings exhibit characteristic stiffness gradients, with greatest stiffness at the base and leading edge and lowest stiffness at the tip of the trailing edge. Additionally, wings are subject to both aerodynamic and structural damping. The sensory consequences of stiffness gradients and damping are unknown. Here we examine how both the nonuniform stiffness profile of a wing and its damping impacts sensory performance, using finite elements analysis combined with sensor placement optimization approaches. We show that wings with nonuniform stiffness exhibit several advantages over uniform stiffness wings, resulting in higher accuracy of rotation detection and lower sensitivity to the placement of sensors on the wing. Moreover, we show that higher damping generally improves the accuracy with which body rotations can be detected. These results contribute to our understanding of the evolution of the nonuniform stiffness patterns in insect wings, as well as suggest design principles for robotic systems.

## 1 Introduction

To execute precise movements, animals and robotic systems alike rely on sensory feedback conveying information about their environment and their own body conformation. Mechanosensory information processing is especially important in movement control. From the capacity to detect pressure and vibration for manipulation tasks to the crucial role of propriception in locomotion, living systems deploy diverse sensory mechanisms for encoding mechanical information. In all of these systems, sensors are embedded in complex, three-dimensional structures whose geometry (size and shape) and material properties (stiffness, damping, density) transform forces into deformations. The information encoded by mechanosensors is therefore not only determined by properties of the sensors themselves but is fundamentally shaped by properties of the structure in which those sensors are embedded.

The interplay between structure and sensing is particularly important in the case of flight. As insects beat their wings, the wings undergo dramatic deformations due to a combination of inertial and aerodynamic forces. Information about these deformations is encoded by a population of strain-sensitive structures sparsely arrayed over the surface of the wing [1], and information about wing deformation is used to guide behavior during flight [2]. The characteristics of wing deformations are determined both by how the animal moves its wings as well as by the structural properties of the wings. While the impacts of wing structure on aerodynamic performance are well studied in both biological and engineered systems [3, 4, 5, 6, 7, 8, 9, 10, 11, 12, 13], the impacts on sensory performance are relatively unknown.

Previous work on the sensory functions of wings has been limited by relying on highly idealized wing models [8, 14, 15]. These models are unable to capture the rich morphological characteristics of real insect wings or the diverse array of structural properties one might consider in designing an engineered system, such as complex shape and nonuniform material properties. These properties will necessarily impact wing bending, and therefore will impact the sensory signals encoded by mechanosensors in the wing. Properties such as shape and stiffness significantly impact wing deformations, the resulting aerodynamic forces, and an animal’s behavioral performance [16, 17, 5, 18]. Models based on the finite element method (FEM) support examination of more realistic wing features, such as wing geometry, venation patterns, damping, and nonuniform stiffness [16, 19, 8, 20, 21, 22]. While these models have revealed an interesting advantage of nonuniform stiffness for thrust and lift generation, they have not yet been used to examine sensing. Nonuniform structural properties will result in more complex spatiotemporal patterns of wing strain and could significantly impact sensing performance and optimal sensing strategies.

In this work, we employ computational approaches to examine how sensing performance depends on wing structure, namely the stiffness gradient of the wing and its damping properties. We develop a series of flapping wing models using the finite elements method, simulating wings with a range of stiffness and damping properties. For each of these wings, we then encode information about strain in a population of optimized neural-inspired sensors and evaluate whether inertial perturbations, which induce subtle changes in wing bending, can be detected via simple decoding mechanisms. We show that nonuniform stiffness improves sensing performance across a range of other model properties. Further, increased damping generally increases sensing performance and reduces the need to fine-tune other properties of the wing to maximize performance. Thus, in addition to the advantages of nonuniform stiffness in the production of flight forces, we demonstrate here additional advantages to sensing. These results provide an inroad to our understanding of the joint evolution of sensing and actuation mediated by the structural dynamics of wings and additionally suggest possible strategies for improved design of engineered flight systems.

## 2 Methods

Our goal is to assess how sensing performance depends on the structural properties of flapping wings. To examine this, we first simulate the complex spatiotemporal patterns of strain that arise in flexible, flapping wings using a finite element analysis (FEA) model. We then encode this strain by simulating neural-inspired spiking sensors in a dense grid over the surface of the wing. These simulations are modeled after strain sensors found on the wings of flying insects. Finally, we use a previously established optimization method [23] to identify the locations of a small population of sensors from which information about wing rotation can best be read out. We perform this series of steps for multiple wings with different structural properties — namely different average stiffness, stiffness gradient, and damping ratio — to determine how sensing accuracy and optimal sensor locations depend on these properties.

### 2.1 Finite element model

An FEA model is created in COMSOL Multiphysics^®^ 5.6 software to study the dynamics of a flapping wing and the effects of various structural properties on the complex spatiotemporal strain distribution on the wing. The finite element model is developed for a geometrically simplified wing that shares several features with the wings of the hawkmoth *Manduca sexta* (e.g., size and flapping frequency). Unlike previous work, which relied on analytical models based on Euler-Lagrange equations, finite element models provide the flexibility to study a variety of complex properties of real insect wings, such as irregular shapes, inhomogeneous material properties, and anisotropies [16, 22].

Here we focus on a few specific aspects of wing structure and model the wing as a single 25 mm × 50 mm rectangular plate with a thickness of 127 μm, as shown in Fig. 1. The thickness is considered to be uniform throughout the plate and the effect of venation is neglected in the simulations. The material properties used in the FEA model are a density of 1180 kg/m^3^ and Poisson’s ratio of 0.35. The material is considered to be linearly elastic. The impact of flexural stiffness on the strain patterns is studied by varying Young’s modulus (*E*) of the plate. The range of explored values for the modulus spans two orders of magnitudes around 3 GPa, corresponding to the average of experimentally measured Young’s modulus for *Manduca sexta* [24].

**Figure 1:**
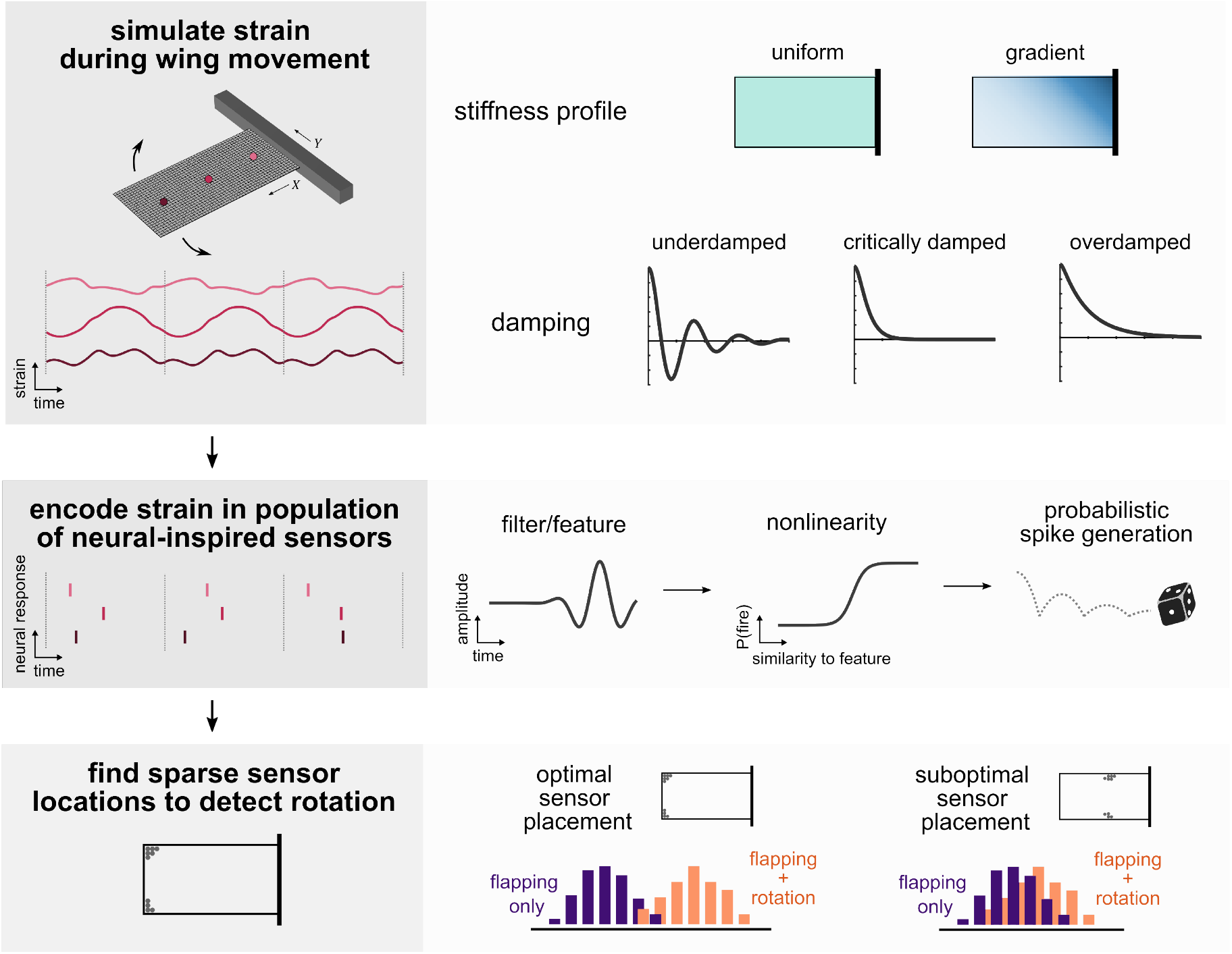
Assessing sensing performance on wings with different structural properties. *Top:* We use a finite element model to simulate strain in flapping wings with different structural properties. Wings may be uniformly stiff (green) or follow a stiffness gradient (shaded blue), from stiffest at a corner of the wing base to most compliant at the opposite corner on the wing tip. Wings may be either overdamped, critically damped, or underdamped. The output of a simulation for a given wing is strain over time at each location on the wing (e.g., light, medium, and dark pink lines). *Middle:* Strain at each location is encoded in a neural- inspired spiking sensor. The strain signal at each location is convolved with a filter, representing a temporal feature of strain the sensor is most sensitive to. Filtered strain is then passed through a sigmoidal function to determine the probability of firing a spike, from which spike are probabilistically generated. Colored vertical lines indicate spike times at locations with corresponding colors above. *Bottom:* Optimization methods are used to identify a sparse set of sensor locations (gray dots) from which information about rotation can best be decoded. Optimal sensor placement will result in greater separation between responses in flapping wings (purple) and responses in wings flapping while undergoing rotation (orange).

Experimental evidence from prior studies suggests that the flexual stiffness of insect wings is not uniform. Specifically, flexual stiffness decays logarithmically, diagonally across the wing from the leading edge base to the trailing edge tip [16]. To understand the effect of this nonuniformity on the capability of the wing to detect rotations, two groups of wings with a (i) uniform, and (ii) nonuniform distribution of Young’s modulus are studied.

In the first group of wings, a constant Young’s modulus (*E_c_*) is assigned throughout the plate. In the second group of wings, a logarithmic decay in flexural stiffness of the flapping wing is prescribed through a spatially variable Young’s modulus *E*_(X,Y)_:

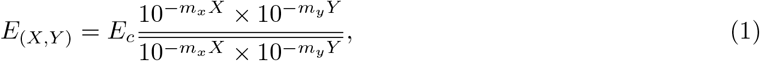

where *X* and *Y* are the positions on the wing surface, and *m_x_* and *m_y_* are logarithmic decline rates along *X* and *Y*, respectively. The overbar in the denominator indicates the mean value of function 10^−*m_x_X*^ × 10^−*m*_*y*_*Y*^ within the wing. Eq. 1 ensures a spatial distribution of stiffness around a mean value of *E_c_*. To achieve two orders of magnitude difference between the minimum and maximum E in the plate (approximately what is observed in hawkmoth wings [16]), a value of 26.67 is applied to the decline rates *m_x_* and *m_y_* in both directions.

Damping is another structural characteristic of the system with significant effects on the dynamic response and the strain patterns generated due to flapping and rotation. Here, viscous damping is used and it is applied isotropically in the form of volumetric forces to the wing. The damping force is defined as *F* = –*cẊ*, where *Ẋ* is the velocity vector at each time instant, and *c* is the damping coefficient of the structure defined as

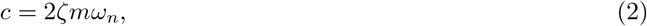

where *ζ* is the damping ratio, *m* is the mass of the wing, and *ω_n_* is the natural frequency of the structure. Values of *ζ* = 0.2,1.0, and 2.0 are chosen to represent a broad range of damping cases, i.e., underdamped, critically damped, and overdamped, respectively. The resonance frequency for the first vibrational mode of the system is calculated using the Eigenfrequency study in COMSOL. The Eigenfrequency analysis is conducted for each stiffness setup and the results are utilized for determining the damping coefficient of the wing. (See Table S1 for selected results from this analysis.)

Second-order Serendipity elements are used to (i) achieve more accurate results compared to first-order elements while (ii) lowering the computational cost compared to the second-order Lagrange elements. To achieve high-quality meshing, a structured quadrilateral mesh is generated on the largest surface of the wing and is swept through the thickness of the wing to create hexahedral meshing in the domain. To determine the size of the mesh, we conducted two sets of convergence analyses: (i) eigenfrequency analysis, and (ii) tip displacement during flapping of the wing (Fig S1). Using 25 × 50 structured hexahedral elements balances the accuracy and the computational cost of simulations by keeping the error lower than 1% for both cases.

Flapping and rotation about different axes are applied through a gyroscopic structure. A constant rotation is applied about a single axis (either roll, pitch, or yaw) at a velocity of 1 rad/s. Using the results from prior studies [25], the flapping pattern of the wing is defined in terms of two harmonic parts: (1) amplitude *A*_1_ = π/12 and frequency *f*_1_ = 25 Hz, and (2) amplitude *A*_2_ = π/60 and frequency *f*_2_ = 50 Hz:

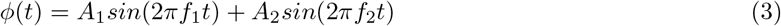

To numerically solve for the wing deformation, we use the implicit method of backward differentiation formula (BDF) with a time step of 2 × 10^-4^ s. To ensure stability of the solver, the maximum order of the BDF is set to 1. In addition, the rotation rate (when applied) is increased from 0 rad/s to its maximum value (1 rad/s) using a ramp function over the course of one flapping cycle to ensure convergence.

### 2.2 Strain encoding in neural-inspired spiking sensors

The local normal strain in the direction of the wing span is then encoded by a dense grid of neural-inspired sensors, from which an optimal subset will be chosen to assess sensing performance, as in previous work [15]. At each node across the 25 x 50 grid of elements (corresponding to every 1 mm), strain is converted to a series of temporally sparse all-or-none sensing events using a linear-nonlinear model, a common model of neural responses [26]. In this model, strain is first convolved with a feature that represents the temporal pattern of strain to which the sensor is most sensitive. The filtered strain is then converted to probability of firing (i.e., triggering a response), *P*(fire), via a nonlinear function, such that greater similarity to the feature results in a higher probability of firing. This function reflects the sensor’s sensitivity to the feature of interest, with high threshold and steep slopes conferring greater selectivity than low thresholds and shallow slopes.

The shapes of the linear filter and nonlinearity are computed from previous electrophysiological recordings of responses in mechanosensors of the wing nerve [27]. The filter f is defined as a decaying sinusoidal function:

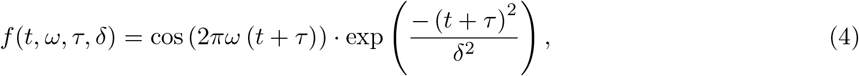

where *ω* is the frequency of the filter, *τ* is the time offset to the peak, and *δ* is the decay time. In this work, 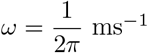, *τ* = 5 ms, and *δ* = 4 ms. The sigmoidal nonlinearity is given by:

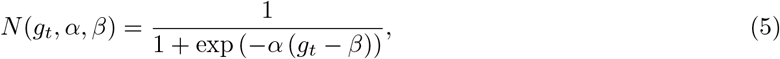

where *g_t_* is the filtered strain at time *t, α* is the slope parameter, and *β* is the threshold parameter, where the function reaches half-maximum. We hold *α* constant at 5 × 10^5^. For results in the main text, the threshold *β* is held constant at 1 × 10^-4^. We also tested the effects of varying the threshold from 1 × 10^-10^ to 1 because the neural threshold substantially impacted results in previous work [15]. In the present study, altering the neural threshold does not alter conclusions. (See Supplementary Information for full results of varying the neural threshold.)

We then generate spikes probabilistically from the output of the linear-nonlinear encoding. The sensor spikes if the probability of firing exceeds a random draw from a standard uniform distribution. We manually impose an absolute refractory period of 15 ms between spikes. This is not intended to represent the actual absolute refractory period of mechanosensors, but rather to empirically match observations from previous experimental work that each sensor fires only 1–2 spikes per wingbeat [28, 29]. For each unique wing and body rotation condition, we generate 100 spiking responses for each sensor. Although the simulated strain is identical at a given location across these repetitions, stochasticity in generating the spiking response results in varied response times at a given location across multiple wingbeats.

### 2.3 Sensor placement optimization

Our objective in sensor placement optimization is to determine the placement of a small number (10) of neural-inspired sensors which can be used to determine whether or not the insect is rotating. We simulate spiking data as described above for two cases: one where a wing is flapping, and one where a wing is flapping and rotating (in the roll, pitch, or yaw axis). For both of these cases, we determine the time to first spike within each wingbeat with 0.1 ms precision and use only this spike timing information to classify the data. For wingbeats where no spike is elicited, we designate the spike time as zero, as in previous work [15]. Data are standardized in this optimization step, but the original (non-standardized) data are used to evaluate accuracy.

To determine optimal sensor locations, we use a previously developed method called *sparse sensor placement optimization for classification* (SSPOC) [30]. This method first uses dimensionality reduction (principal component analysis, in this case) to find a lower-dimensional subspace that captures important features of the data. We then use a linear discriminant analysis (LDA) to find the projection vector *w* that maximally separates the classes of our data in this subspace. Finally, we use elastic net regularization (which linearly combines lasso, or *L*_1_, regularization and ridge, or *L*_2_, regularization) to solve for a sparse set of sensors *s* that can reconstruct the projection vector *w*. We solve:

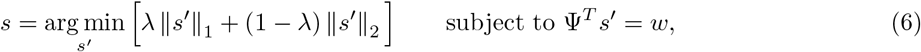

where *s*’ is a vector of sensor weights (*n*x1, with many near-zero entries), *s* is the vector of optimized sensor weights, Ψ is the low-dimensional basis (*n*x*m, m* < *n*), *w* is the projection vector in the low-dimensional subspace (*m*x1), and *λ* determines the balance between lasso and ridge regularization. We set *m* = 3 and λ = 0.9. We use the cvx package to solve this optimization problem (http://cvxr.com/cvx/) [31, 32].

### 2.4 Performance evaluation

90% of the data is randomly selected to be used as a training set for the optimization, and 10% is held out as test data to evaluate accuracy. For straightforward comparison across conditions, we consistently use the top 10 sensors (i.e., sensors with the largest weights in *s*) to assess classification accuracy. In previous work, we find that 10 sensors are typically enough to achieve near-peak accuracy without including a large number of extraneous sensors [15]. This is also a reasonable number of sensors for engineered systems to ensure reduced noise and latency along with higher classification accuracies [33]. LDA is again used to find the best projection vector *w_c_* for the non-standardized test data for only the top 10 sensors. A decision boundary is drawn at the mean of the two condition centroids.

## 3 Results

Our goal is to assess how the structural properties of wings, namely the stiffness gradient and damping, impact an insect’s ability to detect behaviorally relevant perturbations to the wing. For simplicity, we focus on the task of identifying an insect’s body rotation from the strains in its flapping wing. Rotations at realistic speeds produce pertubrations in wing strain that are several orders of magnitude smaller than strain produced by wing flapping [14], making this a challenging detection task.

Examples of the results obtained at various points throughout our process for sensor placement optimization are shown in Figure 1. To evaluate detection accuracy for a wing, we first simulate wing bending as the wing undergoes a periodic flapping motion using the finite element model (Figure 1, *top*). We calculate the time-varying strain at each location on a dense grid along the wing (1 mm spacing; example locations shown in Figure 1, top: dark, medium, and light pink lines). Next, we encode this strain in a population of neural-inspired sensors (Figure 1, *middle*), where sensor properties are chosen to reflect the characteristics of mechanosensitive neurons in insect wings. (See Methods for details.) Finally, we identify an optimal subset of ten sensor locations to detect wing perturbations using the sparse sensor placement optimization for classification (SSPOC) approach (Figure 1, *bottom*) [23].

We repeat this approach for different model wings with varying structural properties to assess the impact of these properties on sensing accuracy. We simulate wings across a range of stiffness values, of both uniform stiffness and logarithmically decaying stiffness, the latter reflecting stiffness gradients observed in insect wings [24]. For each average stiffness and stiffness gradient, we simulate wings that are underdamped, critically damped, and overdamped. We separately assess the ability to detect rotation about the roll, pitch, and yaw axes in each of these wings.

To start, we focus on the impacts of nonuniform stiffness, a common property of insect wings that has been shown to be favorable for aerodynamic performance [19]. Regardless of a wing’s stiffness gradient, strain produced by flapping versus flapping with body rotation are nearly identical, with the strain difference between these two cases approximately three orders of magnitude smaller than the strain produced by flapping alone (Figure 2A,E; wings matched for average stiffness with otherwise identical properties). The spatial profiles of strain over the surface of the wing are different for uniform and gradient stiffness wings, with particularly unique spatial profiles when considering the difference between flapping and flapping with rotation (Figure 2B,F). When sensors are optimized for the uniform stiffness wing, rotation detection accuracy is only slightly above chance performance (Figure 2C,D). However, for the gradient stiffness wing, rotation can be detected by optimally placed sensors with 100% accuracy (Figure 2G,H). Note that spike timing changes caused by the addition of rotation are generally relatively small (e.g., orange and purple spike times in Figure 2G). Optimal sensors are not necessarily found at the locations with the overall greatest strain differences between the two conditions over the course of the wingbeat; rather, locations with large strain differences *at the typical time of spiking* are likely to result in the best ability to distinguish these two cases.

**Figure 2:**
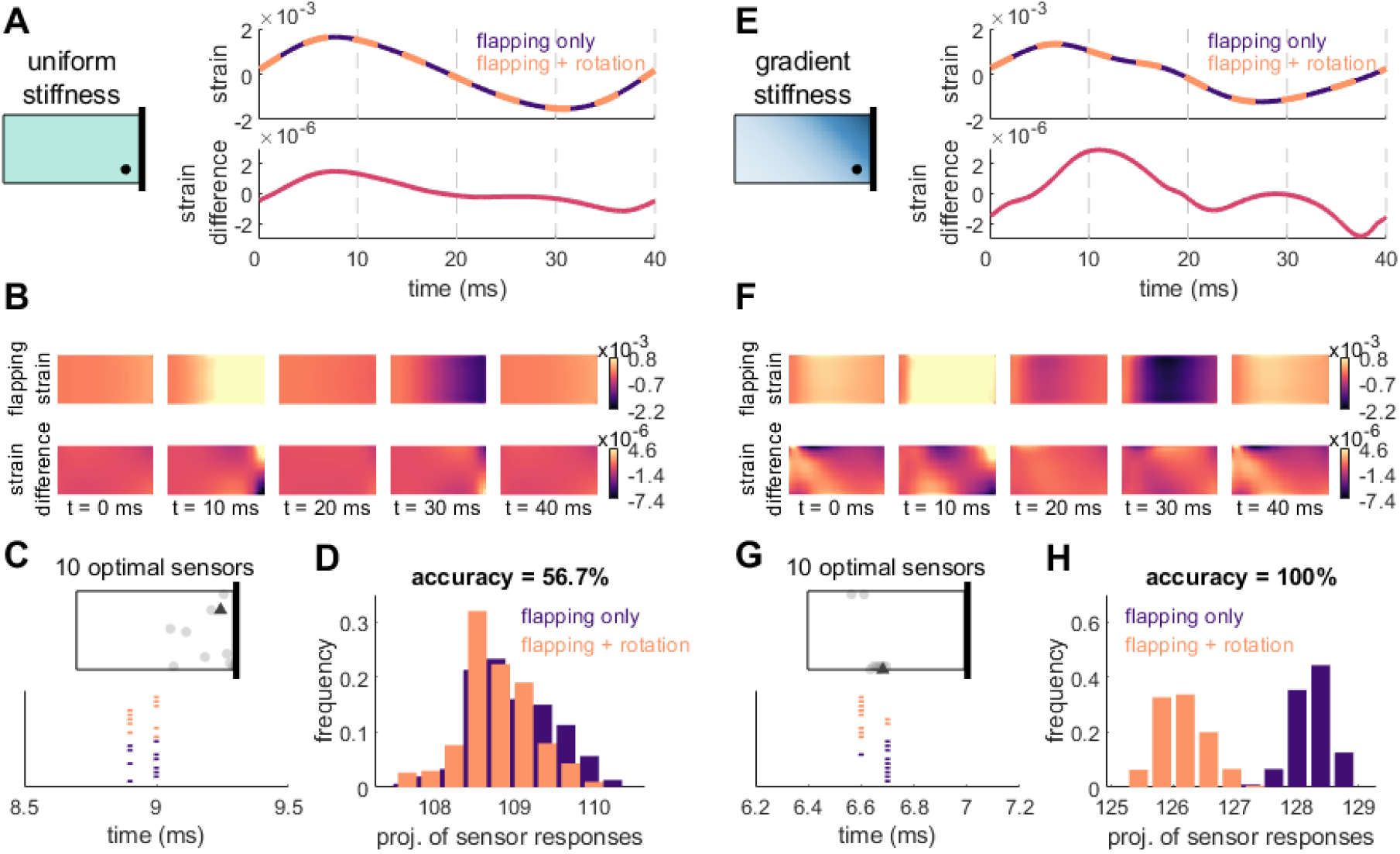
Simulated strain and optimal sensor locations for two example wings, one with uniform stiffness and one with a stiffness gradient. A: *Top:* Strain over the course of a single wingbeat for a uniform stiffness wing that is undergoing flapping only (purple) or flapping along with rotation about the yaw axis (orange). *Bottom:* Strain difference between flapping only and flapping with rotation. B: Top: Spatial patterns of strain at select time points for a flapping wing (without rotation). *Bottom:* Spatial pattern of the strain difference between the flapping only and flapping with rotation cases at select time points. C: Top: Optimal locations for 10 sensors to detect rotation about the yaw axis. *Bottom:* Sensor response times for the single best sensor, indicated by a dark gray triangle in the wing schematic above, on trials when either the wing was only flapping (purple) or flapping while also rotating (orange). D: Projection of the optimal sensor responses onto the vector that maximally separates the two groups. Greater separation between the two distributions leads to greater accuracy in determining whether the wing was rotating. E–H: Same as A–D for a wing with a stiffness gradient, with average stiffness (3.0 GPa) matched to the wing in A.

To examine which mechanisms may underlie the greater accuracy observed for wings with nonuniform stiffness, we conducted a modal analysis of the spatiotemporal strain profiles for each wing. This allowed us to identify differences in spatial patterns of strain and temporal patterns of these spatial modes over the course of each wingbeat for uniform versus gradient stiffness wing (Figure 3A-C). Spatial and temporal modes are similar across both wings for the first three modes, though small spatial asymmetries can be seen for the gradient stiffness wing. Despite the apparent similarity of these modes, their contributions to wing strain allow for clear differences in classification accuracy (Figure 3D). We assessed classification accuracy as a function of the number of modes used to reconstruct wing strain, finding measureable differences in accuracy even for a single mode and maximal differences in accuracy with only three modes. These results demonstrate that the improved performance of nonuniform stiffness wings arises from fundamental differences in wing strain produced by the wing structure, rather than subtle differences in higher-order modes, which may be more influenced by small parameter changes or numerical errors from the finite element method.

**Figure 3:**
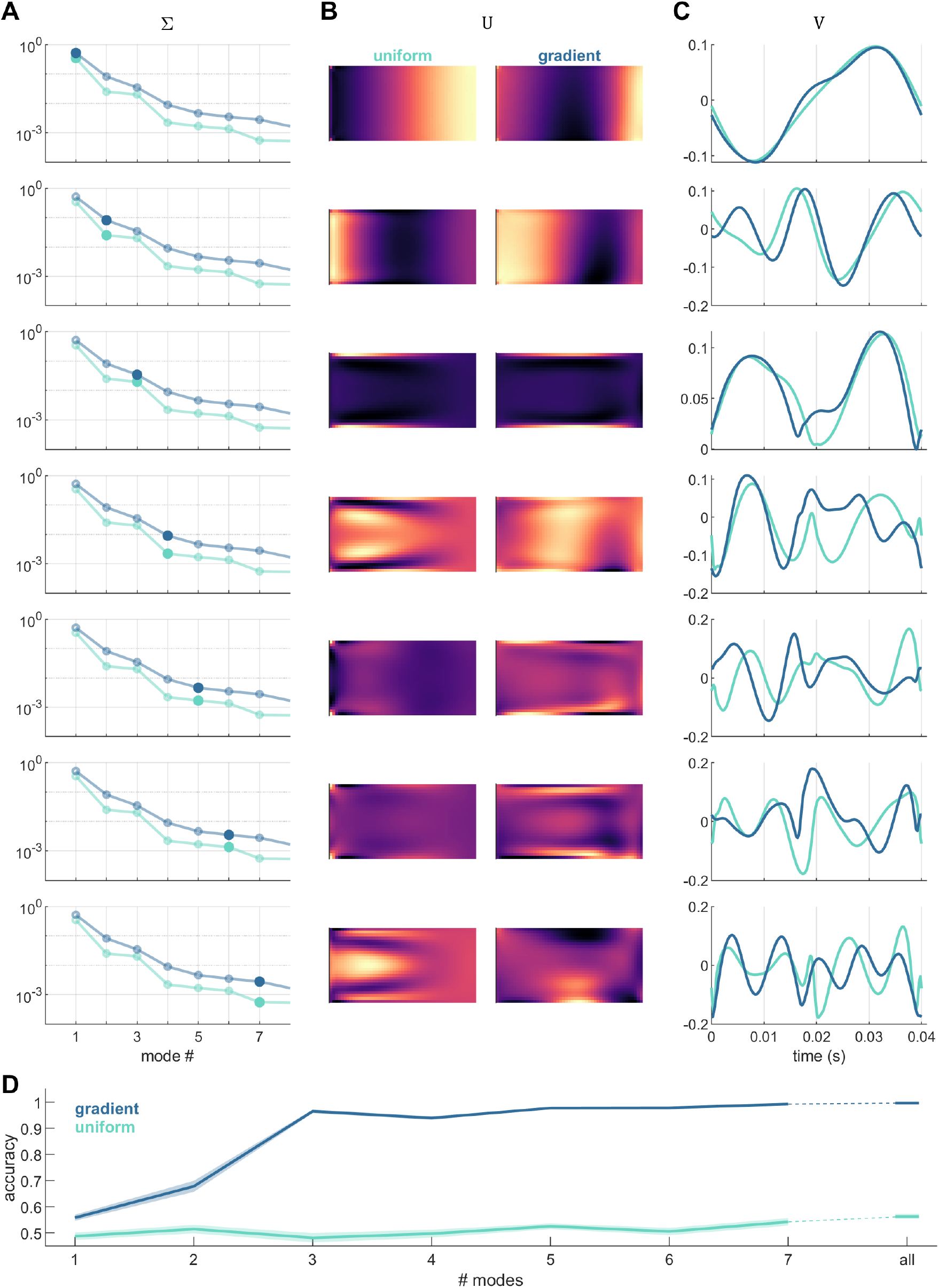
Modal analysis of wing strain shows that few modes are needed to distinguish results in wings with different stiffness gradients. A: Eigenvalue spectra reflecting the relative contributions of each mode to the total strain. B: Spatial strain modes. C: Temporal strain modes. D: Accuracy as a function of the number of modes used to characterize wing strain. Shaded regions indicate SEM over 20 iterations.

We further investigated how the observed changes in wing strain translate to changes in the timing of sensor responses underlying classification. Relatively small changes in mean wing stiffness impact spike timing and can dramatically impact detection accuracy, even when the stiffness gradient, damping ratio, and other properties remain the same. This can even lead to nonmonotonic changes in accuracy as wing stiffness changes, where wings with small differences in stiffness exhibit large differences in accuracy while wings with large differences in stiffness exhibit similar accuracy (Figure 4A). For underdamped wings of uniform stiffness, for example, a very flexible wing may exhibit large differences in spike timing when experiencing flapping only versus flapping with a pitch rotation (Figure 4B). Similar accuracy can be achieved in a wing of moderate stiffness (Figure 4C) or relatively high stiffness (Figure 4D) with very small differences in average spike timing but very precise spike timing. Between two stiffness values that produce high accuracy, an intermediate stiffness may have similarly precise spike times but that occur at nearly identical average times when flapping only compared to flapping with rotation (Figure 4F).

**Figure 4:**
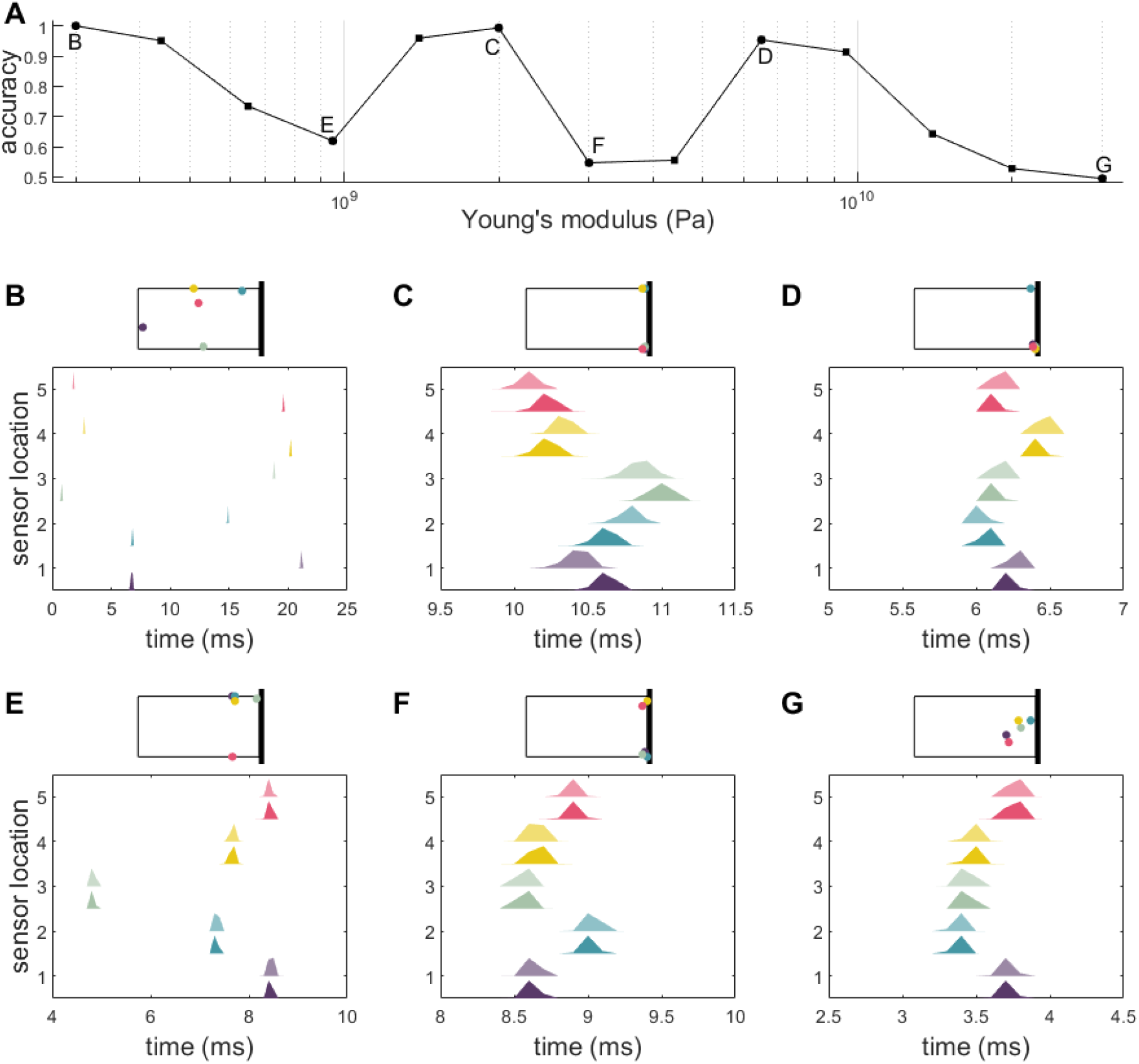
Changes in spike timing underly nonmonotonic changes in accuracy as a function of stiffness. A: Accuracy of pitch detection as a function of wing stiffness for an underdamped wing of uniform stiffness. B–G: Spike timing histograms for top 5 sensors, with locations indicated by dots of corresponding colors in the above wing schematic. (5 shown for simplicity, though 10 sensors used to determine classification accuracy.) For each sensor, dark color indicates spike timing on flapping only wingbeats, while lighter color indicates spike timing on wingbeats with flapping and rotation. Greater differences in spike timing histograms between the two conditions (dark and light colors) results in greater classification accuracy.

We next compare detection accuracy for wings with different stiffness gradients across a range of average stiffness values, different damping ratios, and three different axes of rotation (Figure 5). For wings matched on other characteristics (i.e., mean stiffness and damping), gradient stiffness wings generally outperform uniform stiffness wings. This trend holds true for almost all damping ratios and axes of rotation for almost all average stiffness values tested. Further, wings with higher damping ratios (i.e., overdamped wings) generally result in higher accuracy when other characteristics are held equal. Accuracy typically decreases as average stiffness increases, though the relationship may be nonmonotonic (e.g., pitch detection for a uniform stiffness wing with damping ratio 0.2, also shown in Figure 4). Lower damping ratios tend to produce more irregular relationships between accuracy and average stiffness in addition to generally lower accuracy.

**Figure 5:**
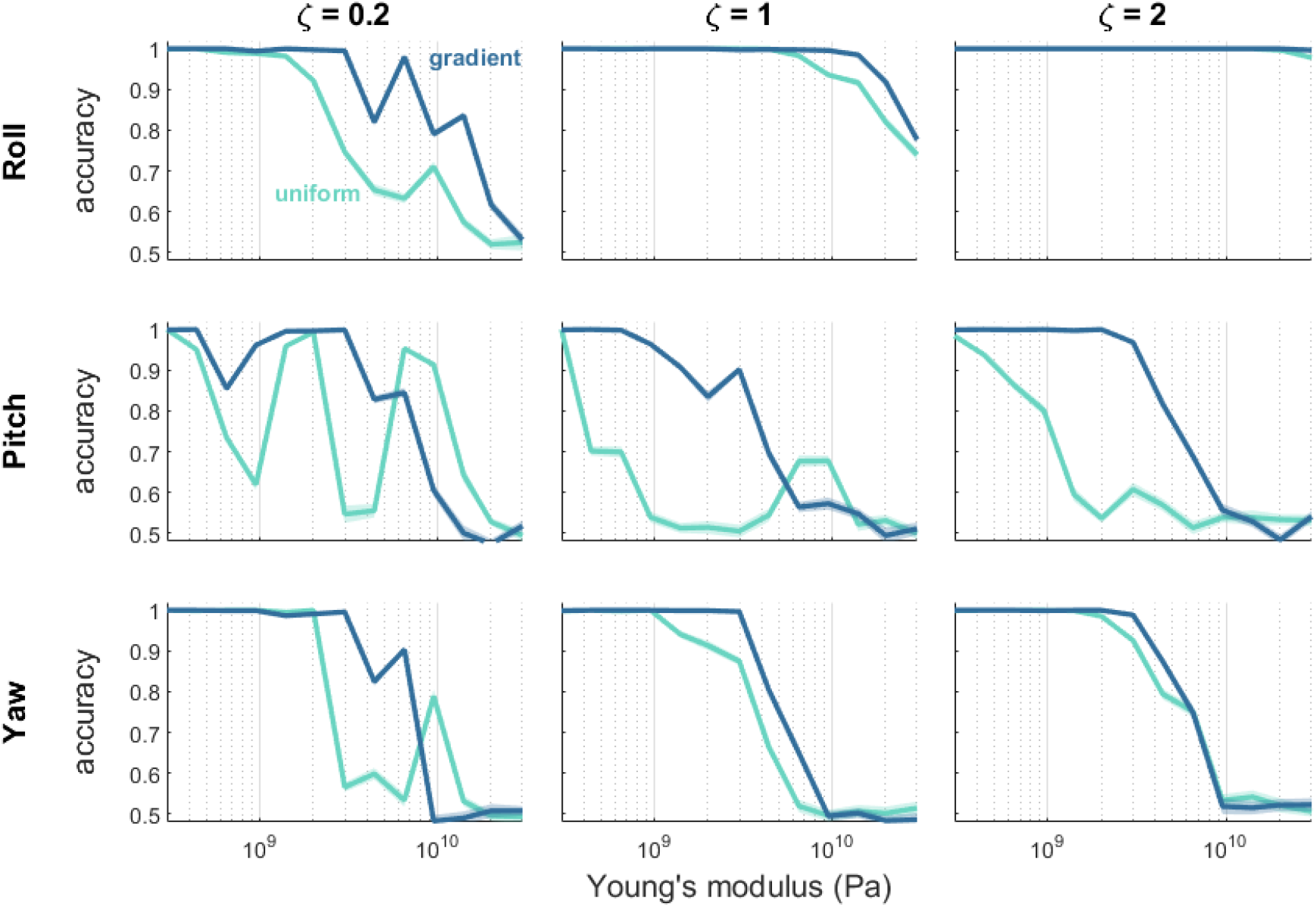
Wings with a stiffness gradient typically result in higher classification accuracy compared to uniform stiffness wings matched for average stiffness. Each panel shows classification accuracy for groups of 10 optimally placed sensors as a function of average wing stiffness. Different subpanels show results for under-, critically, and overdamped cases (columns) and for rotation about different axes (rows). For each condition (damping, axis of rotation, average stiffness, uniform or gradient) sensor optimization was performed for 20 separate iterations to assess classification accuracy. Shaded regions show SEM.

Previous work showed that optimal sensor placement shifted between locations clustered at the wing base and locations clustered at the wing tip, depending on wing average stiffness and the axis of rotation to be detected [15]. However, these observations were limited by the wing model it used, which not only studied wings of uniform stiffness but also restricted the wings to taking on linear combinations of characteristic modal shapes. The direct method in finite element models we employ in the current work, however, is not concerned with mode truncation and allows more complex solutions. Therefore, we next examine how the average spanwise locations of optimally placed sensors depend on features of wing structure (Figure 6). Like previous work on uniform stiffness wings, sensors are typically localized near the wing base across a range of other parameters, though as average stiffness decreases sensors shift distally. On gradient stiffness wings, on the other hand, optimal sensors are found far more distally — generally at intermediate locations on the wing. Note that the stiffness gradient for these wings runs from the leading edge of the wing base (stiffest at the top right corner as depicted) to the trailing edge of the wing tip (most flexible at the bottom left corner), i.e. diagonally. On a given wing, optimal sensors are generally not placed along the line of equal wing stiffness.

**Figure 6:**
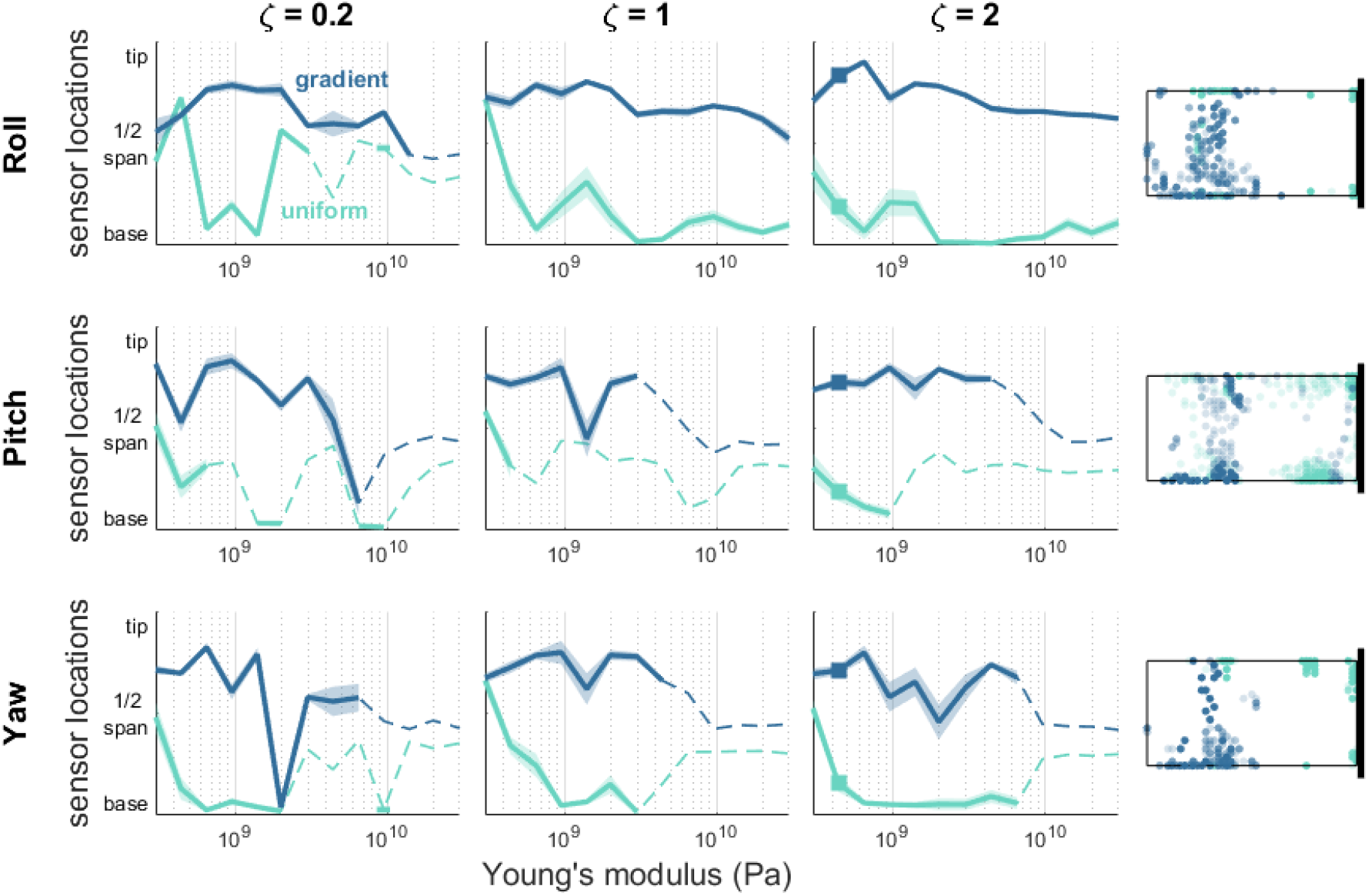
Wings with a stiffness gradient result in optimal sensors placed more distally (towards the wing tip) compared to uniform stiffness wings matched for average stiffness. Average spanwise location (from wing base to wing tip) of 10 optimal sensor locations is shown as a function of average wing stiffness. Shaded regions show SEM. Solid lines show results when average classfication accuracy (Fig. 5) exceeds 70%. Dashed lines show when accuracy falls below this threshold. Note that sensor placement becomes random as classification accuracy approaches chance performance. Right column shows the distribution of optimal sensor locations (10 sensors for each of 20 iterations) for example cases indicated by squares in *ζ*=2 panels.

We have shown that stiffness gradients and overdamped wings generally have sensing advantages across a range of other wing properties and detection tasks for *optimally placed* sensors. However, insects need not have sensory structures optimally positioned, and even on robotic systems, sensor placement may be constrained by other design demands. To examine how closely sensing accuracy depends on opimtal sensor placement, we next consider how much classification accuracy suffers when sensors are placed randomly (Figure 7). We calculate a measure of *placement sensitivity:* the classification accuracy of 10 optimized sensors divided by the accuracy of 10 randomly placed sensors minus 1. This results in values ranging from 0 to 1, where 0 indicates no sensitivity to sensor placement (i.e., random sensors perform equally as well as optimized sensors), and 1 indicates complete sensitivity (i.e., random sensors perform at chance when optimized sensors perform with perfect accuracy). Accuracy in stiffer wings is typically more sensitive to random sensor placement. This is consistent with the general decrease in accuracy observed for these wings: as wings become stiffer, the classification task becomes more difficult, resulting in decreased detection accuracy and increased reliance on particular sensor placement. Although relatively similar across a range of other parameters, placement sensitivity tends to be modestly higher for uniform stiffness wings compared to gradient stiffness wings, indicating that gradient stiffness wings allow for more flexible sensor placement across a range of other wing properties and sensing tasks.

**Figure 7:**
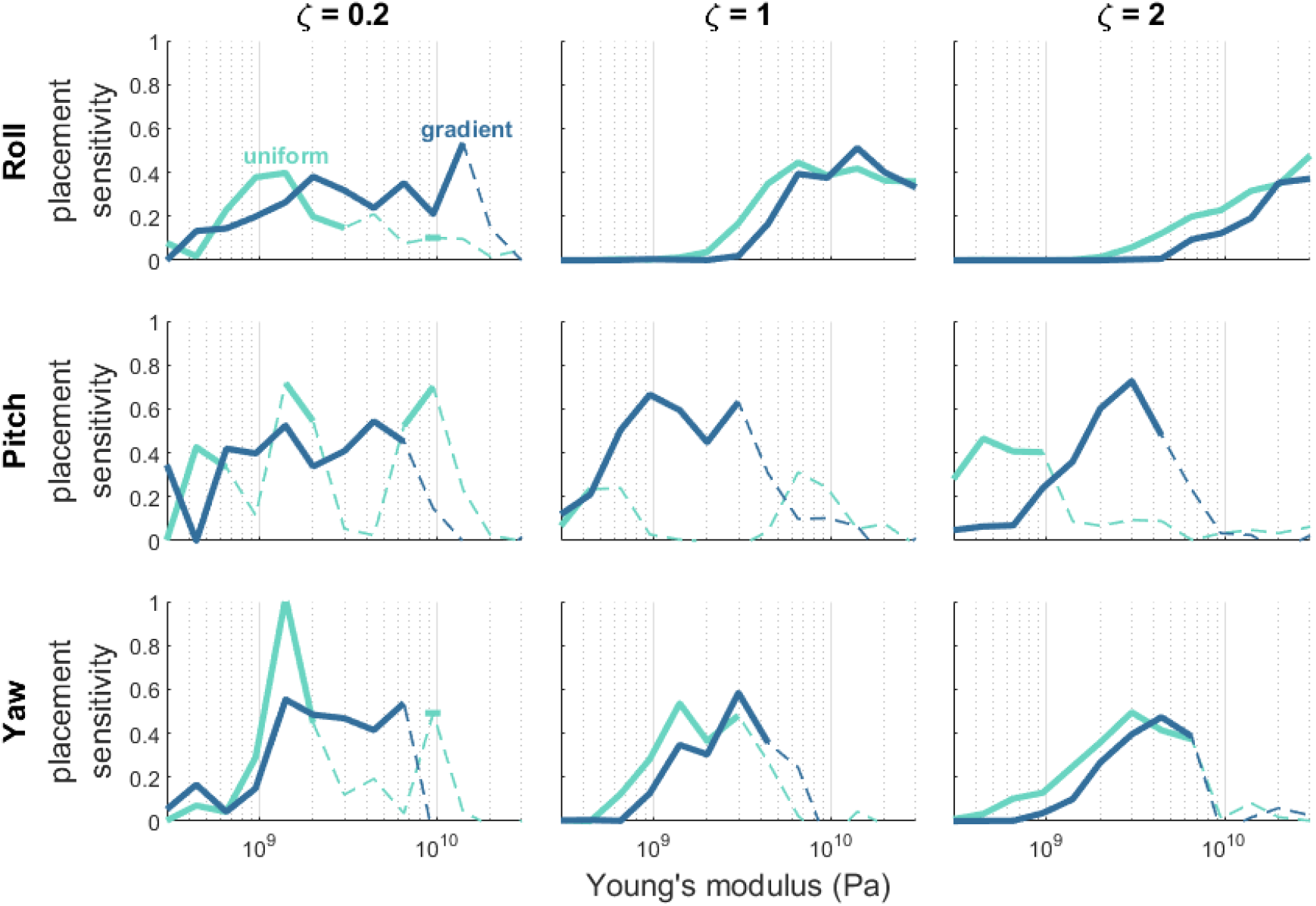
Uniform stiffness wings require more careful sensor placement than wings with a stiffness gradient. Each panel shows classification accuracy of 10 optimized sensors divided by accuracy of 10 randomly placed sensors minus 1 as a function of average stiffness. A value of 1 indicates maximum sensitivity to sensor placement: optimized sensors can achieve perfect accuracy while random sensors perform at chance. A value of 0 indicates no sensitivity to sensor placement: random sensors perform equally as well as optimized sensors. Solid lines show results when average classification accuracy (Fig. 5) exceeds 70%. Dashed lines show when accuracy falls below this threshold.

## 4 Discussion

We use finite element models to examine how structural properties of a wing impact its ability to sense perturbations to body dynamics during flight. Our results point to two general conclusions regarding sensing performance for flaping wings: those with a spatial gradient in stiffness and those with a higher damping ratio enable more accurate detection of body rotations. Wings with a stiffness gradient exhibit higher detection accuracy across a range of average stiffness values, damping ratios, and across rotation axes (Figure 5). Additionally, wings with gradient stiffness are less sensitive to particular sensor placement, with higher detection accuracy for randomly placed sensors relative to that for uniform stiffness wings (Figure 7).

### Implications for insect flight

Insect wings exhibit stiffness gradients from wing base to tip and leading to trailing edge [16]. The aerodynamic consequences of wing stiffness have been studied extensively, with nonuniform stiffness exhibiting a number of advantages, including significant improvements in the production of both lift and thrust forces in flapping wings and flutter-induced drag reduction [19, 34, 7, 35, 36]. Our results suggest that an additional function of this stiffness gradient may be to aid sensing performance, by directly improving accuracy and/or by allowing for more flexible placement of sensors on the wing. Further, gradient stiffness wings exhibit smoother changes in accuracy as average stiffness changes. This may provide robustness to changes in wing stiffness that occur over an insect’s lifespan [37] and may prevent the need for evolutionary fine-tuning of wing stiffness for sensing purposes. Thus, the current work suggests that aerodynamic and sensing needs may present coordinated evolutionary pressures that result in the observed stiffness gradients of insect wings.

We show that increased damping generally improves sensing performance, while also increasing the smoothness of accuracy as a function of wing stiffness. Insect wings have previously been reported to be underdamped [38, 39], although other work has reported that wing veins in particular are overdamped [40]. Our own measurements in *Manduca sexta* wings show a combination of underdamped and overdamped properties (Figure S5). Greater damping may help with flight stability, as wing movements will be less affected by external perturbations (such as gusts or collisions) common during flight. Damping may also avoid significant aerodynamic flutter and wing failure. Yet higher damping may limit the amplitude of camber that could lead to favorable aerodynamics forces. Wings with a combination of overdamped and underdamped properties, as we observe in our own tests, may balance the tradeoffs between these two conditions, though more work is needed to elucidate the consequences of the damping properties observed in insect wings.

### Implications for engineered systems

Much like insect wings, engineered wings for flapping-wing micro aerial vehicles (MAVs) do not have uniform structural properties, and the methods proposed in this work are likely relevant for similar classification tasks like quickly sensing disturbances and rotations in these small vehicles (e.g., [41, 42, 43, 44, 45]). Even though these vehicles can often be equipped with an inertial sensor that can measure rotation rates, wing deformation has been predicted to precede body rotation and could provide an earlier warning of unexpected rotations [46]. Beyond flapping flight, this sensing and sensor placement approach may be useful in other engineered systems in which event timing can change when disturbances are applied. Legged robots are another example of cyclic systems that need to quickly classify and react to disturbances (e.g., [47]); an unexpected rotation could alter the pressure on the robot foot within a gait cycle.

Given that the classification approach in this paper depends on spike timing within a cycle, the work in this paper is particularly relevant to systems with cyclic behaviors. However, the ability to match a sensing system to the synthetic structure and perturbations of interest can likely be extended to non-cyclic engineered systems. Large arrays of sensors have previously been designed for fixed-wing micro aerial vehicles (MAVs) to gather data on wing deformation during flight with the goal of classifying disturbances before the MAV’s inertial sensor can detect movement [33]. Sensing disturbances like gusts that can deform fixed wings may be facilitated by material choices and stiffness gradients throughout the wing. Similarly, work in the field of soft robotics has shown how useful graded and anisotropic material properties can be for actuation and locomotion [48], but these variations in material properties may also facilitate a robot’s ability to sense unanticipated contact or other disturbances.

Finally, this work suggests an entirely new approach toward sensing than is currently used in engineered systems. The vast majority of engineered systems utilize sensors that provide an analog or digital value when sampled. Instead, as shown in Figure 2, our approach uses differences in event timing (e.g., a probablistic crossing of a threshold) to distinguish rotations that would otherwise be exceedingly challenging to sense with conventional strain sensors. Even the simplest off-the-shelf microcontrollers can distinguish digital events down to sub-microsecond resolution, and recent work has shown several ‘switch’-based sensors that may be utilized in a sensing framework as proposed in this paper (e.g., strain [49], acoustic [50], flow [51]). This approach may provide a step toward low-latency and event-based sensing in engineered systems.

### Future directions

In the current work, we employ finite element models, which allow great flexibility to investigate the effects of complex wing structures. This increased scope of wing shapes and material properties that we can explore, however, comes with the potential for numerical error and instability. We conducted two separate convergence studies to mitigate potential numerical issues (Figure S1). In this study, we retained the highly simplified rectangular wing shape of earlier models, but allowed for more complex spatial patterns of stiffness. However, we are acutely aware that wing morphology — namely, shape and venation pattern — will have significant impacts on the patterns of spatiotemporal strain induced by wing flapping, and will therefore have significant impacts on sensing accuracy and optimal sensor placement as well. Future work should therefore address the effects of different wing morphologies on sensing. Recent work has provided new quantitative tools that bring together data on wing shape and other aspects of wing structure in hundreds of different species [52]. Combining this rich data set with the framework established here can be utilized in future work to study the effects of more realistic wing shape on sensing performance.

This work does not address the potential impacts of aerodynamic forces on sensing in wings. This simplifying assumption is supported by previous work showing that aerodynamic forces have minimal impact on wing bending relative to inertial-elastic forces for *Manduca sexta* [53]. However, aerodynamic forces may more significantly impact bending for smaller wings. Additionally, even relatively small changes in wing bending may be behaviorally significant if they improve an animal’s ability to sense relevant features of wing bending.

Several additional simplifying assumptions were made about how sensors encode information about wing strain and how information from sensors is decoded. For example, sensors encode information about spanwise strain only. Previous work showed that this resulted in better performance than sensors that encode chordwise strain [15]. Additionally, this previous work showed that sensing performance was not improved with more complicated decoding schemes than that used in this work (linear discriminant analysis). As ongoing work uncovers more detail about which features of wing bending biological mechanosensors on insect wings are sensitive to, additional theoretical study may reveal the significance of these response properties.

Previous work on sensing during insect flight has largely focused on the halteres, the modified hindwings of flies that are generally considered to serve as gyroscopic sensors of Coriolis forces [54, 55, 56]. In the current work, we draw on previous studies showing a key sensory function for wings [2, 27] and similarly consider how wings may serve this function, detecting small changes in strain induced by constant-velocity rotations. Yet for an insect in flight, sensing transients — induced by gusts or changes in direction, for example — may be even more behaviorally relevant. The transient changes in wing strain induced by these perturbations are much larger than those induced by constant rotation, and thus likely present an easier sensing task. This may nevertheless be a valuable area of future research due to its likely behavioral relevance.

The structure of insect wings is presumably shaped by many potentially competing demands acting on their evolution. Factors likely include force generation for flight, sensing, mating display, and predator avoidance, among others [57, 58, 59, 60]. Perhaps one of the most promising, but also challenging, areas of future study will investigate the interplay between the many competing evolutionary pressures shaping the evolution of insect wings and flight control.

## 5 Funding

This work was supported by the Air Force Office of Scientific Research [FA9550-18-1-0114: BWB; FA9550- 19-1-0386: BWB], the Washington Research Foundation [AIW], and the eScience Institute at the University of Washington [AIW], with additional prior support from the AFOSR [FA9550-14-1-0398: TD].

## Supplementary Information

### Convergence of finite element simulations

In order to assess how many elements were needed to achieve stable simulations of flapping wings, we conducted a series of convergence studies to analyze simulation error as a function of the number of mesh elements (Figure S1). We opted to use 1,250 elements (25 x 50 grid), which achieved very low simulation error (<0.5%) without excessively long runtimes. We conducted two sets of convergences analyses: (i) Eigenfrequency analysis, and (ii) displacement of the flapping wing tip. In the eigenfrequencey model, we track the effect of reducing mesh size by recording the first natural frequency of the wing. For the flapping wing model, we measure the maximum vertical deflection of the wing tip during a wingbeat to analyze the impact of mesh convergence on the accuracy of solutions.

**Figure S1:**
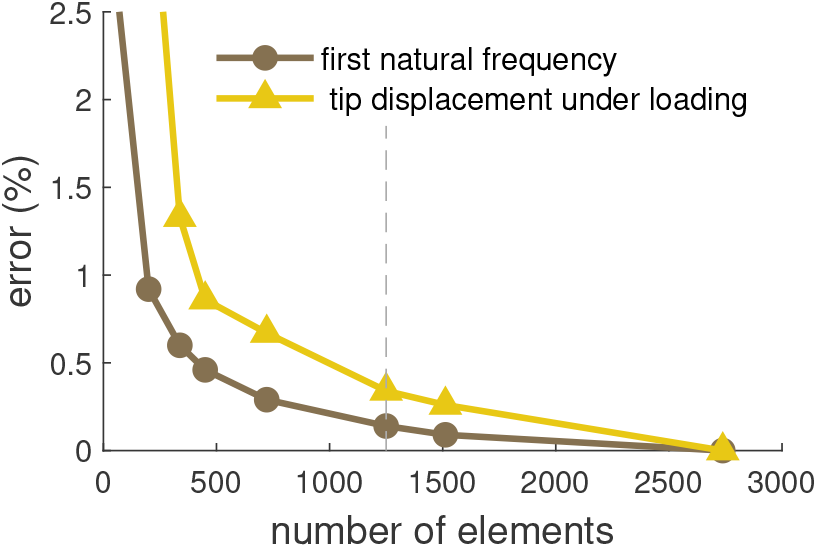
Simulation error as a function of number of mesh elements. The results of two seperate mesh convergence studies are shown. The first natural frequency from the Eigenfrequency analysis of the wing (brown) and the maximum tip displacement of the wing from the flapping wing model (yellow) are recorded to inspect the solution accuracy. A greater number of elements reduces error but also requires longer runtimes.

Additionally, for a subset of wings we ran longer simulations to ensure that our results did not depend on small wingbeat-to-wingbeat changes in simulated strain due to numerical stability. We focused on rotation detection about the pitch axis because these conditions produced the most complex relationship between accuracy and wing stiffness, particularly for underdamped wings. Longer simulations (0.68 ms) resulted in minimal changes in detection accuracy compared to shorter simulations (0.44 ms) and did not affect the overall trends observed (Figure S2).

**Figure S2:**
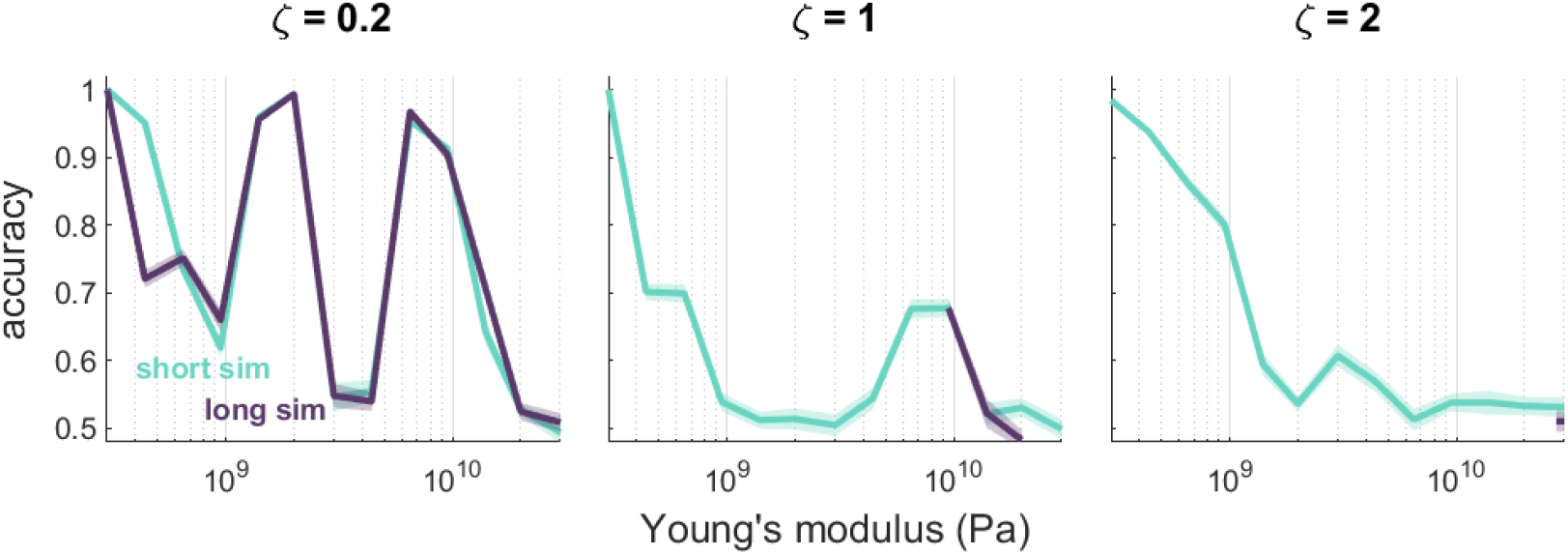
Detection accuracy of rotation about the pitch axis for underdamped (left), critically damped (center), and overdamped (right) wings of uniform stiffness. Results are similar for short simulations (teal, same center row in Figure 5) and longer simulations (purple). Long simulations were run for only a subset of stiffness values for critically and overdamped wings.

**Table S1:**
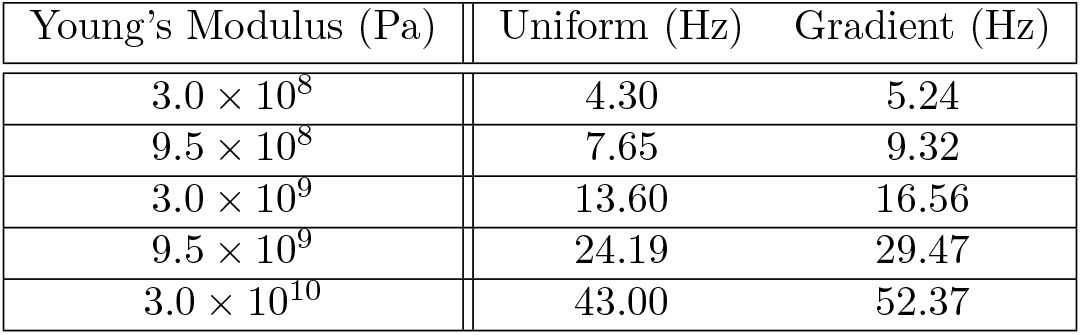
The first natural frequency (*ω_n_*) of uniform and nonuniform wing structures.

| Young's Modulus (Pa) | Uniform (Hz) | Gradient (Hz) |
| --- | --- | --- |
| $3.0 \times 10^8$ | 4.30 | 5.24 |
| $9.5 \times 10^8$ | 7.65 | 9.32 |
| $3.0 \times 10^9$ | 13.60 | 16.56 |
| $9.5 \times 10^9$ | 24.19 | 29.47 |
| $3.0 \times 10^{10}$ | 43.00 | 52.37 |

### Eigenfrequency analysis

The resonance frequency for the first vibrational mode of the system is calculated using the eigenfrequncy study in COMSOL. We conduct the eigenfrequncy analysis for each stiffness setup and use the results for determining the damping coefficient of the wing using Eq. 2 from the manuscript. The full breakdown of the first natural frequencies are presented in Table S1.

### Varying sensor threshold

We examined the effects of varying sensor threshold to ensure that our results did not depend on the specific value chosen. Results reported in the main text use a threshold value of 1 × 10^-4^, though classification accuracy is similar for lower thresholds across a range of wings (uniform: Figure S3; gradient: Figure S4). At higher thresholds, accuracy is generally near chance because these thresholds are sufficiently high to result in little to no sensor response.

**Figure S3:**
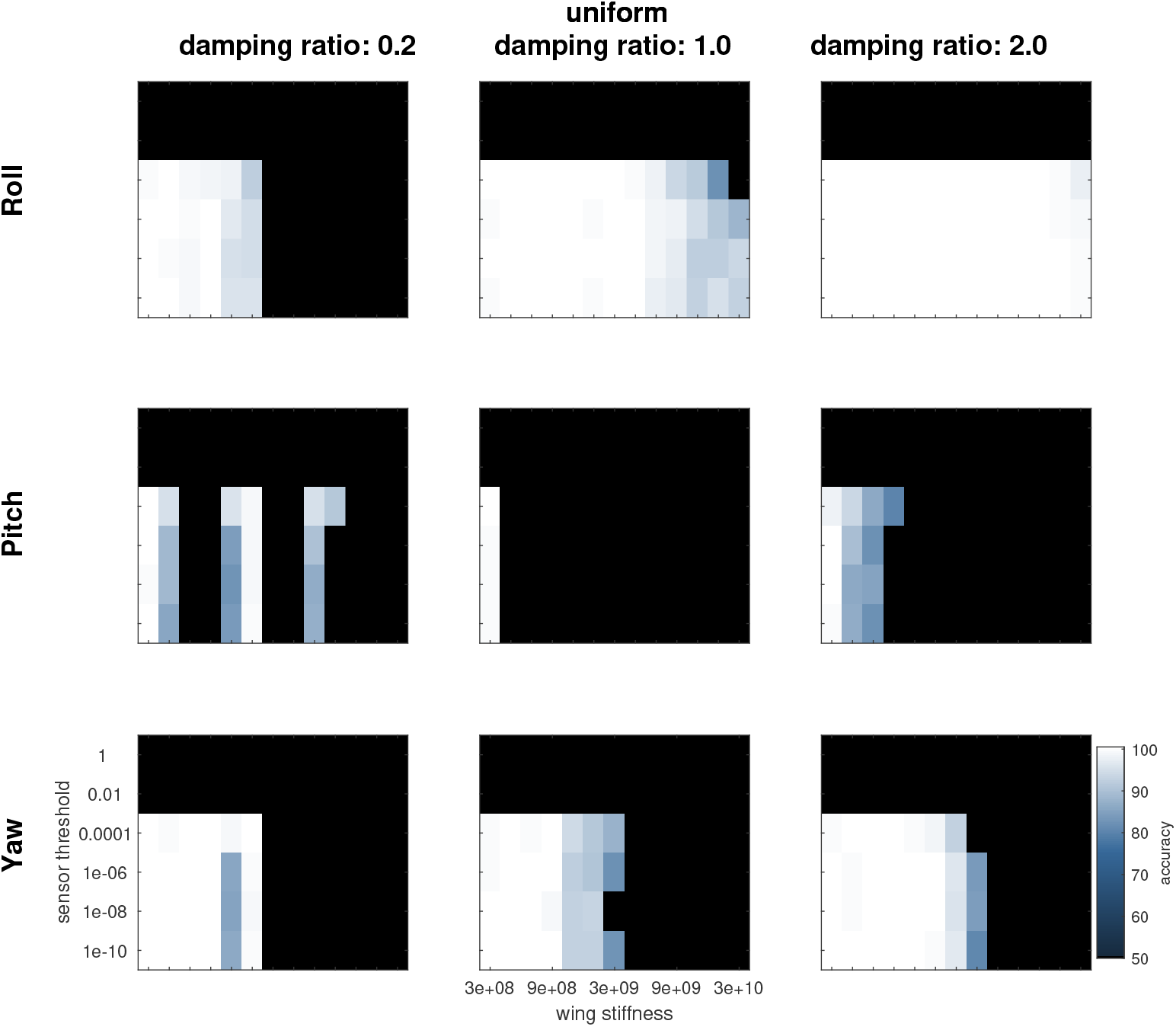
Classification accuracy for uniform stiffness wings as a function of sensor threshold and wing stiffness for each axis of rotation (roll, pitch, yaw) and wings with different damping ratios (0.2, 1, and 2).

**Figure S4:**
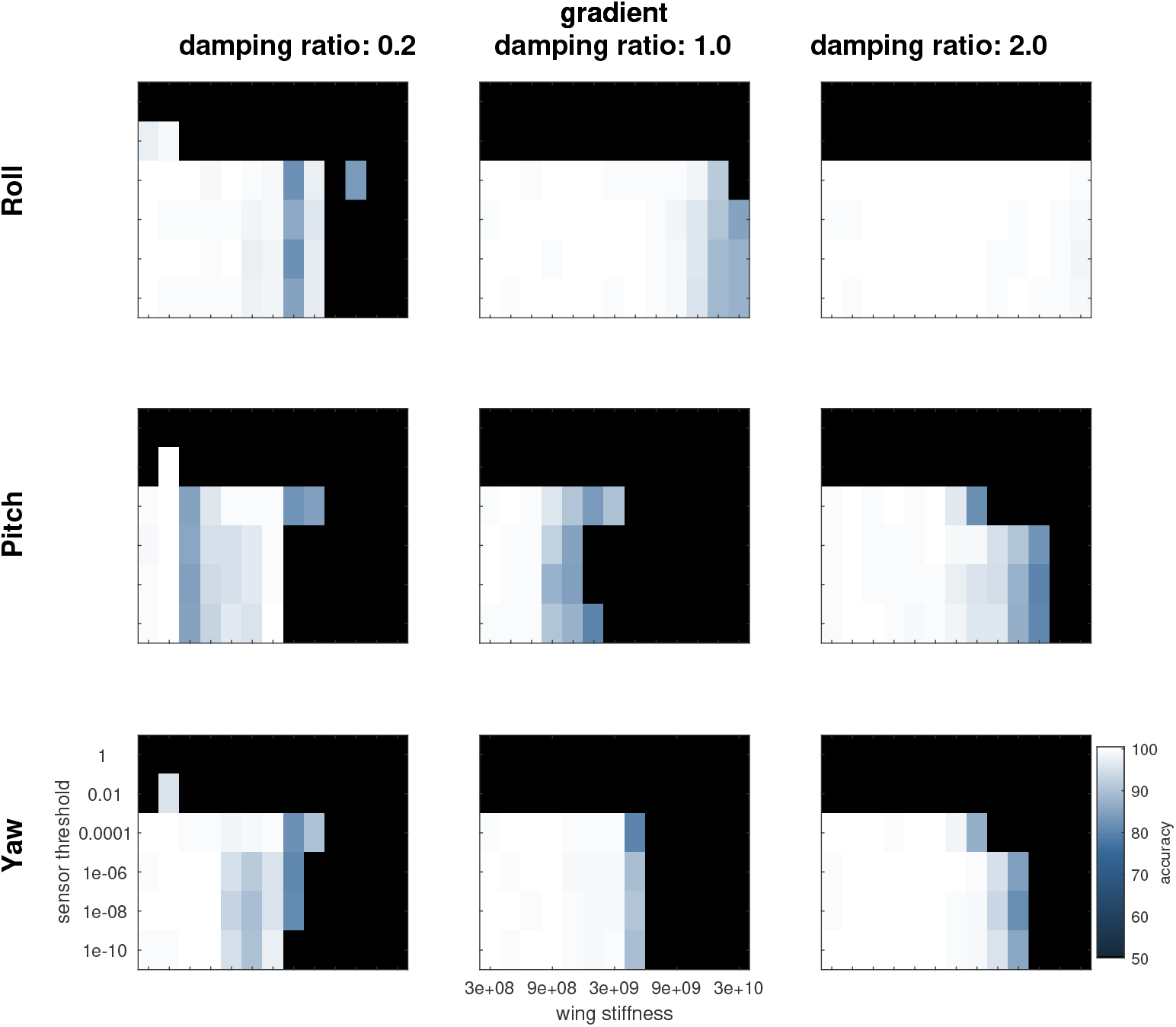
Classification accuracy for nonuniform stiffness wings as a function of sensor threshold and average wing stiffness for each axis of rotation and wings with different damping ratios.

### Experimental measurement of damping in *Manduca sexta* wings

To experimentally determine the damping properties of *Manduca sexta* wings, we used a laser displacement sensor to measure displacement near the wing tip when the wing is released from a deflected state and allowed to passively return to baseline. One characteristic example is shown in Figure S5, in which the wing was deflected in the dorsal direction and then released. The response shows a mixture of underdamped and overdamped characteristics.

**Figure S5:**
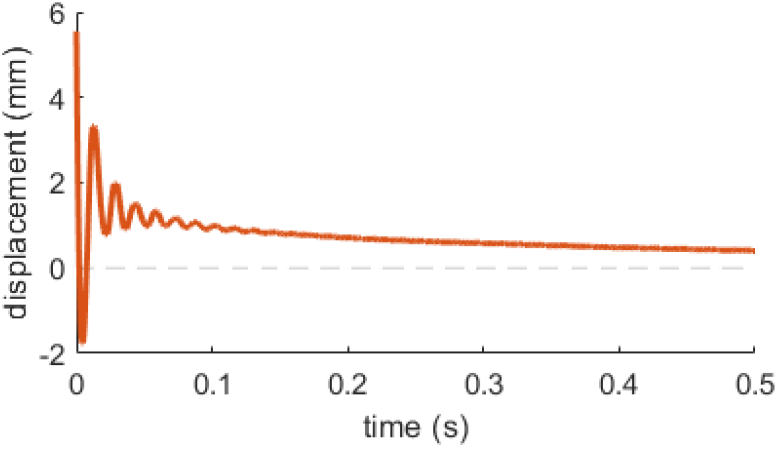
Measured displacement when a wing is released from a deflection and passively returns to baseline.

## Notes

### Competing Interest Statement

The authors have declared no competing interest.

